## Supplemental Material for "Multiparametric quantitative phase imaging for real-time, single cell, drug screening in breast cancer"

**The PDF file includes:**

Supplementary Tables, S1 and S2
Supplementary Figures 1-16
Legends for Movies M1-M10
Description of Supplementary data files

**Other Supplementary Material for this manuscript includes the following:**

Movies M1-M10
Supplementary data files

| Table S1. Cell lines and receptor status | |
|---|---|
| **Cell line** | **Receptor status** |
| MCF-7 | ER+/PR+/HER2- |
| MDA-MB-231 | ER-/PR-/HER2- |
| BT-474 | ER-/PR+/HER2+ |

| Table S2. List of compounds | | |
|---|---|---|
| **Drug name** | **Abbreviation** | **Target** |
| 5-Fluorouracil | 5FU | DNA-RNA synthesis |
| Carboplatin | carb | DNA synthesis |
| Docetaxel | doc | Microtubule |
| Doxorubicin | dox | DNA topoisomerase II |
| Fulvestrant | fulv | Estrogen receptor |
| Lapatinib | lap | EGRF/HER2 |
| Palbociclib | palb | CDK4/6 |
| 4-hydroxy-Tamoxifen | 4HT | Estrogen receptor |
| Vinblastine | vin | Microtubule |

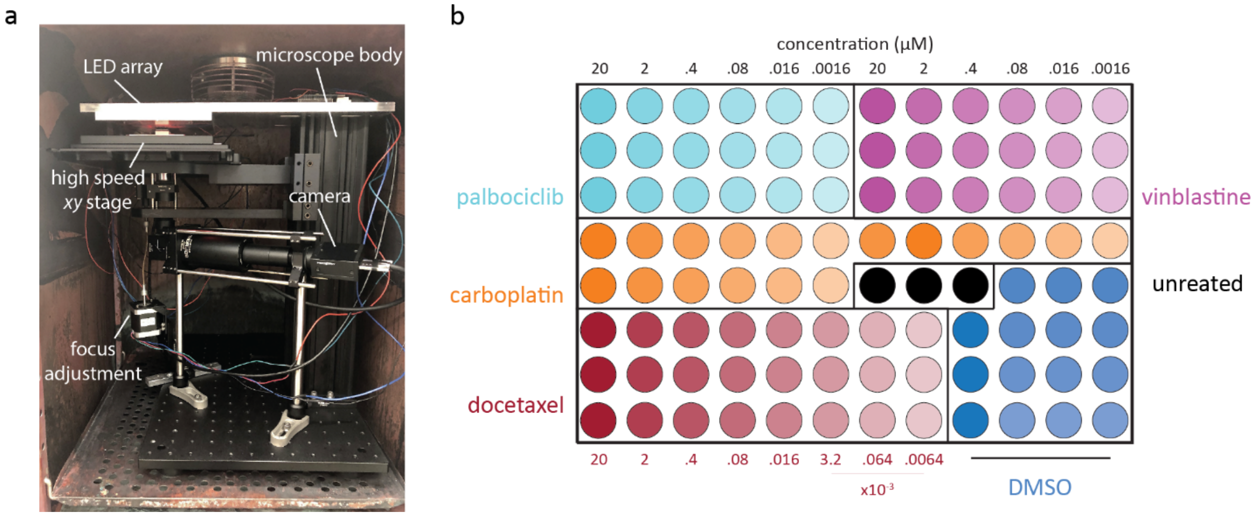


**Supplementary Figure S1:** (a) Photograph shows implementation of DPC QPI microscope placed inside a cell culture incubator for temperature and $CO_2$ control throughout the duration of the experiment. (b) Plate map showing how therapies for drug panel B are organized. Docetaxel uses an expanded range of concentrations to capture the typically low $EC_{50}$s for this drug, noted in maroon below the panel.

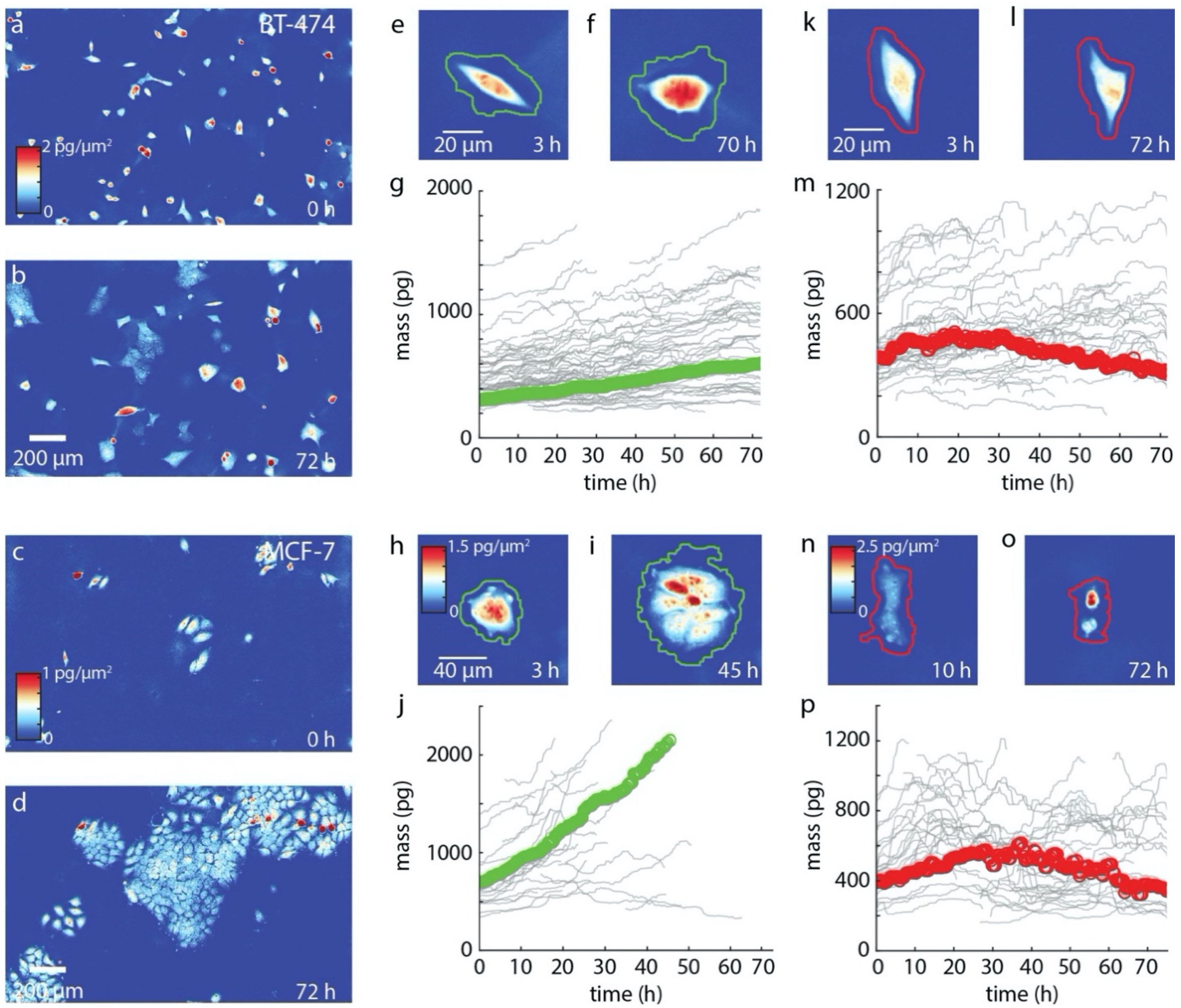


**Supplementary Figure S2: QPI measures the drug sensitivity and temporal dynamics of cell clusters.** (a) Field of view of BT-474 cells treated with DMSO at the beginning of a 72 h experiment (b) and at the end of the experiment where they have grown into clusters. (c) An example of a well containing MCF7 cells at the beginning of the experiment from the DMSO control. (d) By the end of the experiment, the MCF7 cells have grown to fill most of the field of view. (e) Cluster of 3 BT-474 cells from the DMSO solvent control that grow into (f) a larger cluster of 6 cells that are segmented together (green) to measure the growth of the cluster. (g) The individual mass measurements for this cluster are plotted showing robust growth. (h) A small cluster of MCF7 cells treated with DMSO at the beginning of the experiment, (i) grows into a larger cluster by the 45 h timepoint. (j) Measuring the mass of this cluster demonstrates robust growth. (k) An example of a small cluster of BT-474 cells exposed to 2 µM of doxorubicin at the beginning of the experiment. (l) At the end of the experiment, the cluster has not grown noticeably in size. (m) The clusters mass is tracked over time shows the cluster is losing mass. (n) However, MCF7 cells exposed to 20 µM fulvestrant at the 10 h (o) appear to shrink in size over the course of the experiment. (p) Plotting the mass over time for this small cluster, shows that they grow slowly at the beginning and end the experiment with a lower mass than they started with.

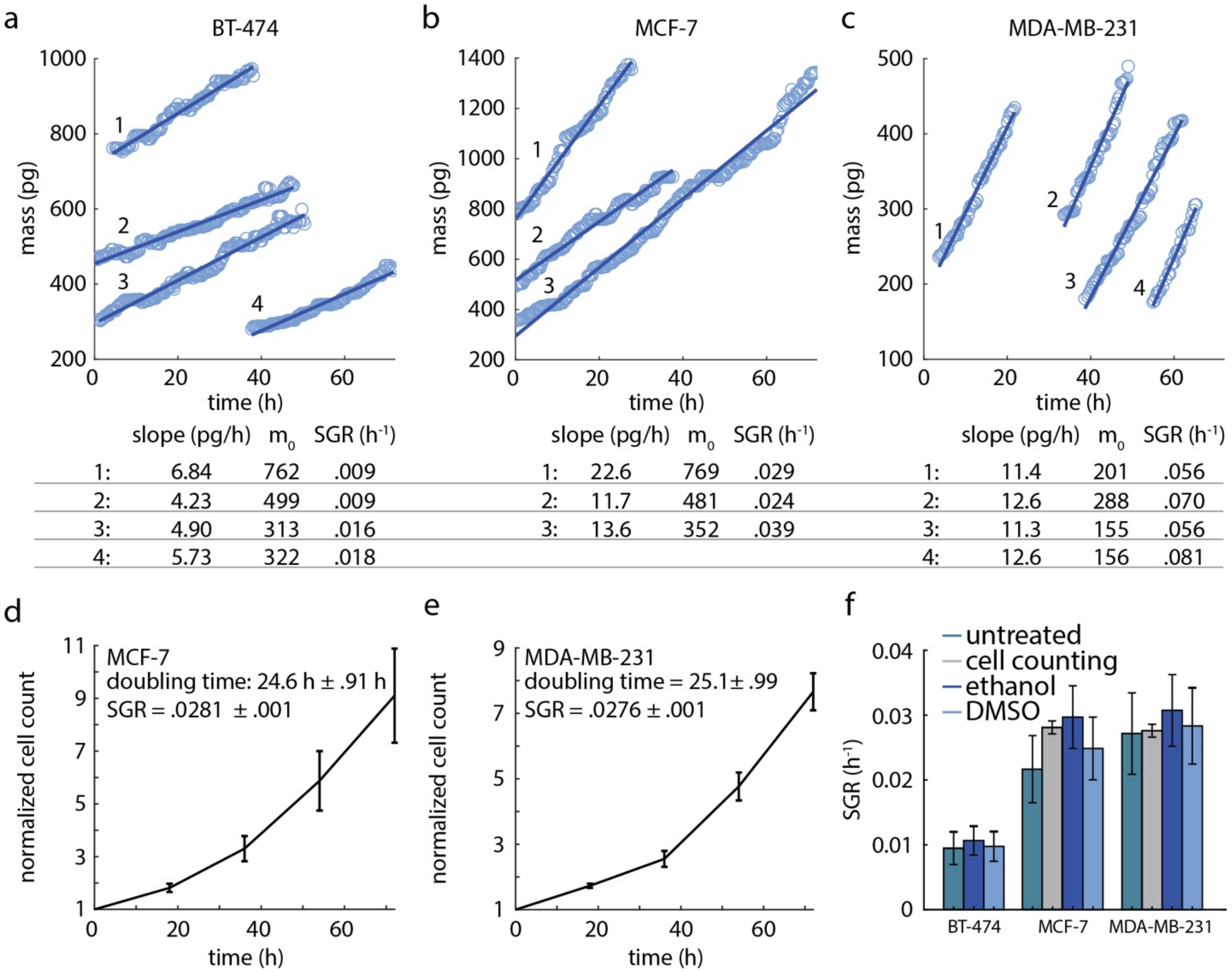


**Supplementary Figures S3: Linear relationship between mass and growth.** (a-c) Mass over time plots for BT474, MCF7, and MDA-MB-231 cells shows a linear fit to measure the growth rate of individual cells. (d) Proliferation measurements of MCF7 cells treated with ethanol shows a doubling time of approximately 24.6 h, which corresponds to an exponential growth constant of 0.0281 $h^{-1}$. (e) Proliferation measurements of ethanol treated MDA-MB-231 cells shows a doubling time of 25.1 h, which corresponds to an exponential growth constant of 0.0276 $h^{-1}$. (f) Growth rate measured using cell counting shows similar results to growth rate measured using QPI.

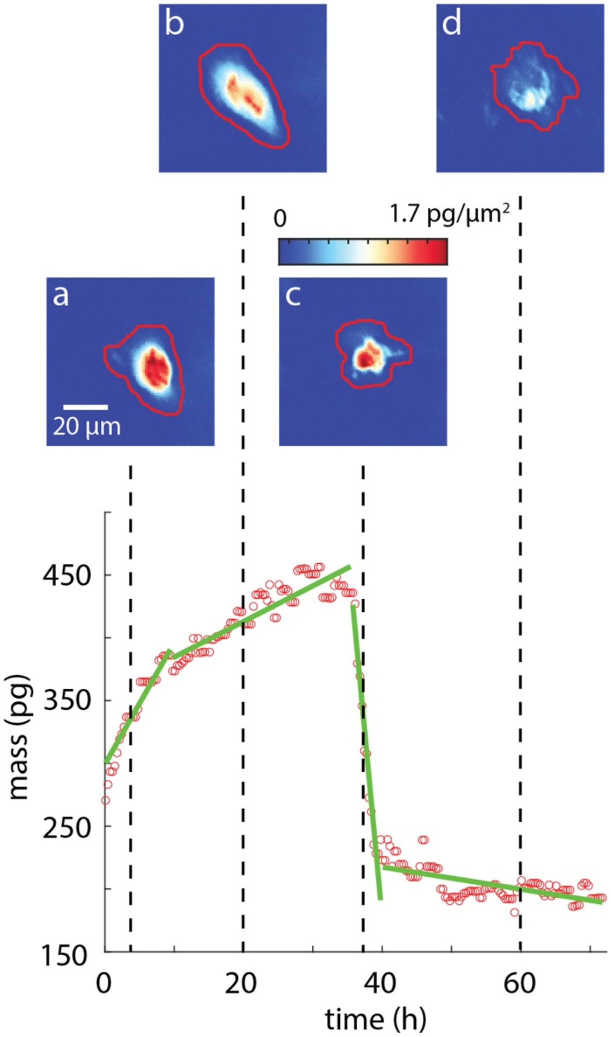


**Supplementary Figure S4: Temporal dynamics of drug response for a single MDA-MB-231 cell during exposure to 2 μM doxorubicin.** (a) Cell initially exhibits robust growth (SGR = 0.0358 $h^{-1}$). (b) After about 13 h of imaging the growth rate of the cell abruptly declines (SGR =0.0105 $h^{-1}$) for the following 19 h. (c) The cell then rapidly loses mass (SGR = -0.217 $h^{-1}$) at approximately 40 h post-exposure during the active stages of cell death. (d) The cell then continues losing mass (SGR = -0.0033 $h^{-1}$) throughout the rest of the experiment.

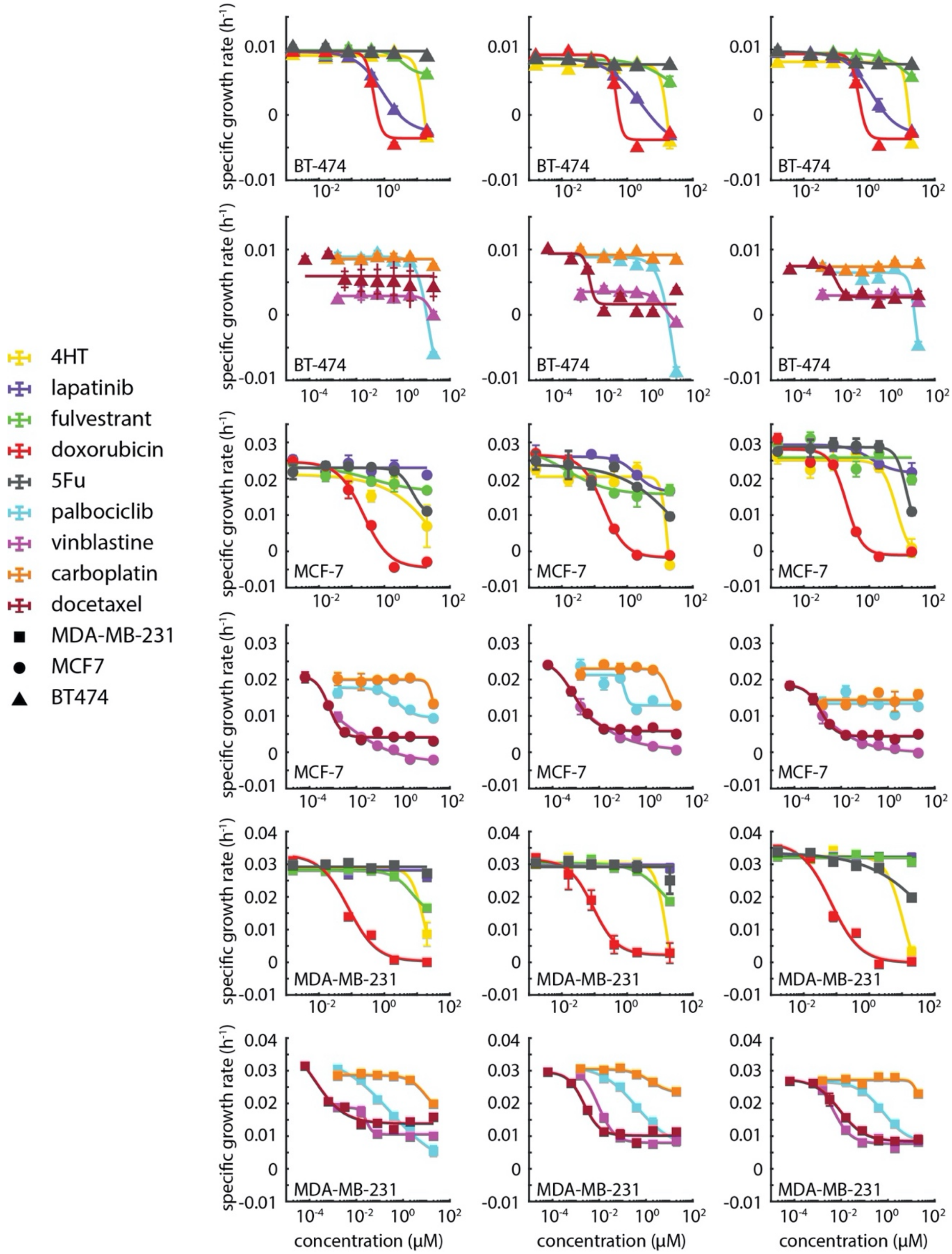


**Supplementary Figure S5: Dose response measurements for all cell lines and drug combinations.** Dose response plots show mean specific growth rate as a function of concentration. A 4-parameter Hill curve is fit to each dose response plot to extract the depth of response and $EC_{50}$ for each cell/drug pair. A flat line represents no response due to failure of the F-test to reject the null hypothesis that the logistic fit is indistinguishable from a flat line.

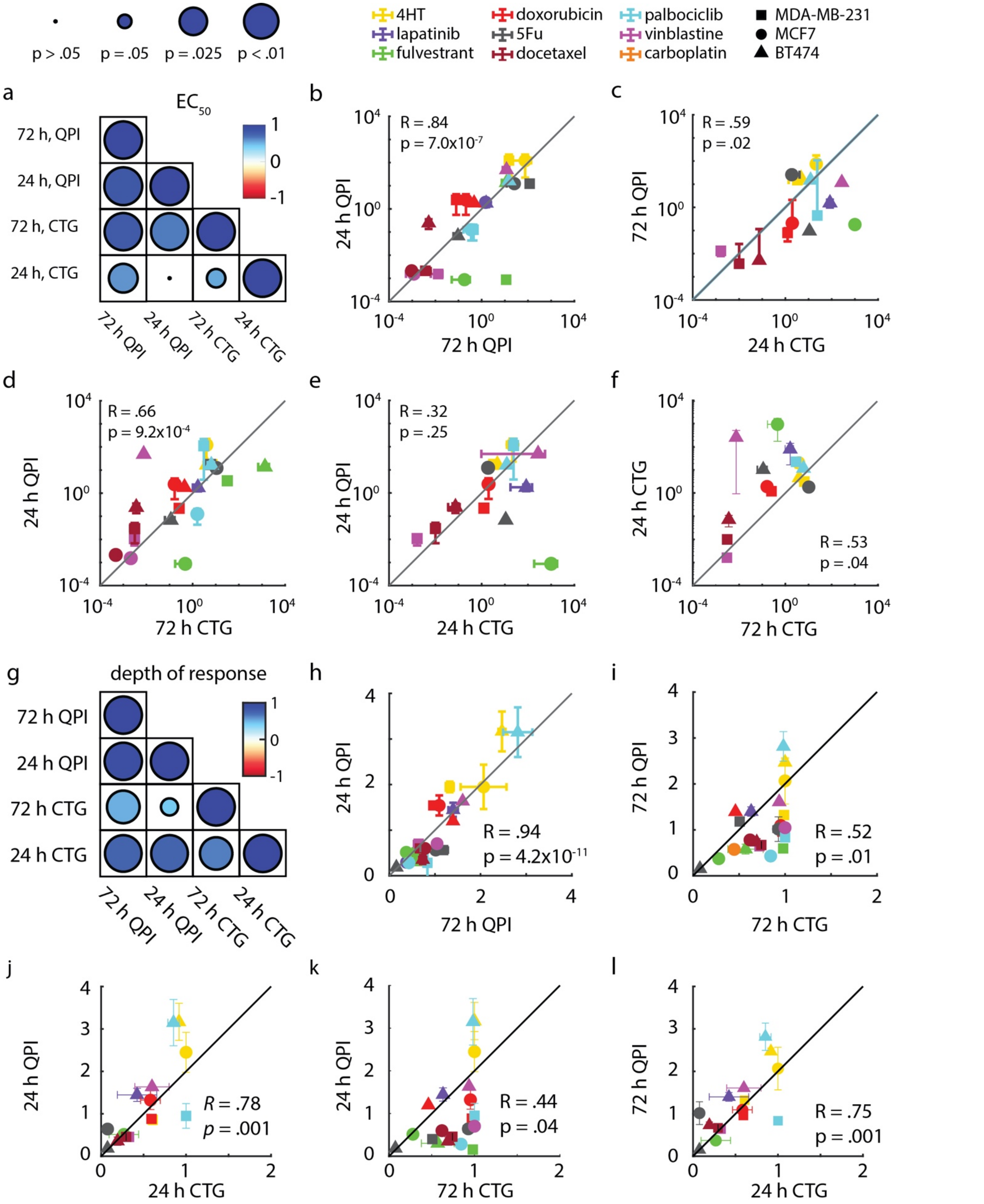
p > .05
p = .05
p = .025
p < .01
4HT
lapatinib
fulvestrant
doxorubicin
5Fu
docetaxel
palbociclib
vinblastine
carboplatin
MDA-MB-231
MCF7
BT474
a
EC50
72 h, QPI
24 h, QPI
72 h, CTG
24 h, CTG
72 h QPI
24 h QPI
72 h CTG
24 h CTG
b
R = .84
p = 7.0x10-7
c
R = .59
p = .02
d
R = .66
p = 9.2x10-4
e
R = .32
p = .25
f
R = .53
p = .04
g
depth of response
h
R = .94
p = 4.2x10-11
i
R = .52
p = .01
j
R = .78
p = .001
k
R = .44
p = .04
l
R = .75
p = .001

**Supplementary Figure S6: 24 h QPI data predicts results in 72 h CTG and QPI experiments.** (a) Correlation map of $EC_{50}$ for both QPI and CTG experiments. Color indicates correlation (blue high, red low) and symbol size indicates *p*-value. (b) $EC_{50}$ values determined from 72 h QPI and from the first 24 h of QPI data are strongly correlated ($R$ = .84, $p = 7\text{x}10^{-7}$). (c) $EC_{50}$ values from 24 h QPI and 72 h CTG are strongly correlated ($R$ = .66, $p = 9.2\text{x}10^{-4}$). (d) $EC_{50}$ values measured in 72 h using QPI and 24 h CTG are moderately correlated ($R$ = .59, $p$ = .02). (e) $EC_{50}$ values obtained using 24 h of QPI data and 24 h CTG are not well correlated ($R$ = .32, $p$ = .25). (f) $EC_{50}$ values measured using 24 h CTG and 72 h CTG are moderately correlated ($R$ = .53, $p$ = .04). (g) Correlation matrix of DoR measured using both QPI and CTG experiments. Color indicates correlation (blue high, red low) and symbol size indicates *p*-value. (h) DoR values measured in first 24 h of QPI and 72 h QPI are strongly correlated ($R$ = .71, $p = 1.4\text{x}10^{-4}$). (i) DoR determined using 72 h QPI and 24 h CTG are not well correlated ($R$ = .48, $p$ = .08). (j) The DoR values determined in the first 24 h of QPI and 24 h CTG are strongly correlated ($R$ = .71, $p = 3.0\text{x}10^{-3}$). (k) DoR measurements from the first 24 h of QPI and 72 h CTG are moderately correlated ($R$ = .5, $p$ = .02). (l) DoR measurements obtained using 72 h of QPI and 72 h CTG are moderately correlated ($R$ = .5, $p$ = .01). The gray line is y = x. The black line is y = 2x. Error bars show SEM.

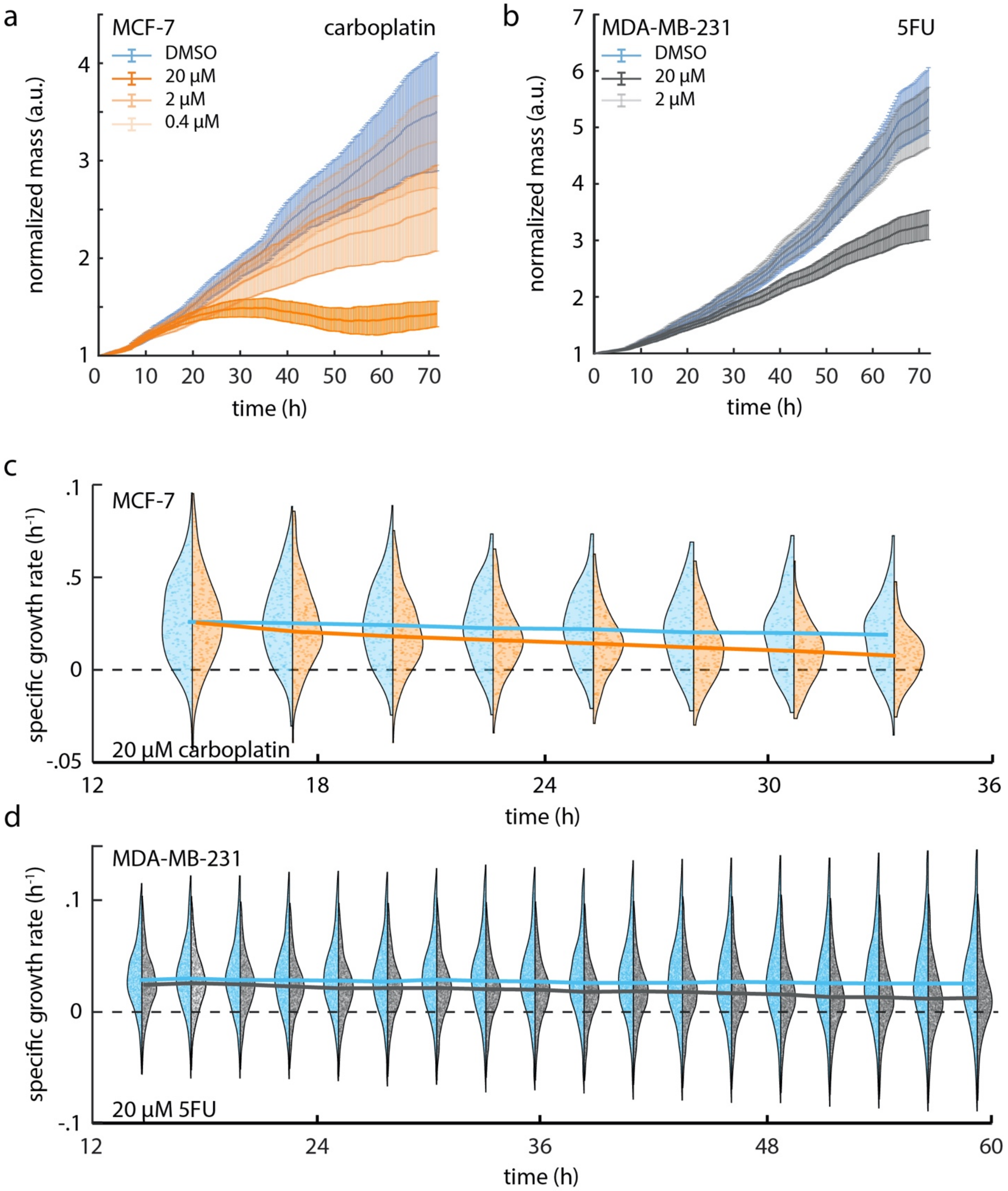


**Supplementary Figure S7: Dynamic response plots for MCF7 and MDA-MB-231.** (a) Average mass plotted against time, for carboplatin treated MCF7 cells. Mass is normalized by the initial mass of each cell in the population. Dark orange represents 20 μM carboplatin treated cells, and lighter colors show the same plot for cells treated with lower concentrations. (b) Average mass plotted against time for 5-fluorouracil (5FU) treated MDA-MB-231 cells. Mass is normalized by the initial mass of each cell in the population. Dark gray shows the normalized mass of cells treated with 20 µM 5FU and lighter gray shows cells treated with 2 µM 5FU. Error bars show SEM. (c) BT-474 response to 20 µM carboplatin (orange) versus DMSO control (blue) in 12 h bins shows an initially similar distribution that slowly begins to deviate over time. Solid lines represent the median of the distribution as a function of time. Dashed line shows SGR of zero. Individual data points shown within population distributions. (d) MDA-MB-231 response to 20 µM 5FU (gray) versus DMSO control (blue) in 12 h bins shows an initially similar distribution that slowly begins to deviate over time. Solid lines represent the median of the distribution as a function of time. Dashed line shows SGR of zero. Individual data points shown within population distributions.

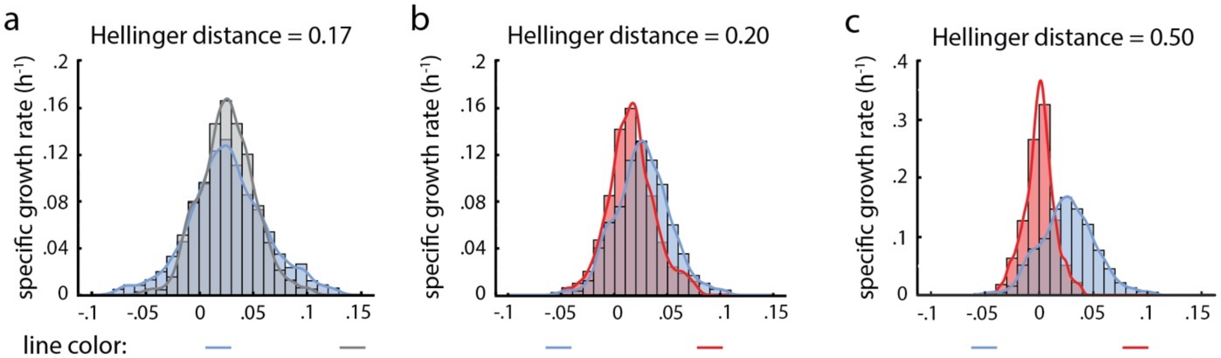


| | | | | | | |
|---|---|---|---|---|---|---|
| treatment: | DMSO | DMSO | DMSO | doxorubicin | DMSO | doxorubicin |
| concentration: | control | control | control | 0.08 μM | control | 0.4 μM |
| time: | 28 h | 58 h | 28 h | 28 h | 28 h | 28 h |
| mean: | 0.0262 $h^{-1}$ | 0.0261 $h^{-1}$ | 0.0262 $h^{-1}$ | 0.0129 $h^{-1}$ | 0.0262 $h^{-1}$ | 0.00 $h^{-1}$ |
| SD: | 0.0256 $h^{-1}$ | 0.0359 $h^{-1}$ | 0.0256 $h^{-1}$ | 0.0239 $h^{-1}$ | 0.0256 $h^{-1}$ | 0.0138 $h^{-1}$ |

**Supplementary Figure S8: Hellinger distance used to determine the statistical distance between drug treated and control treated groups.** (a) The specific growth rate distributions for DMSO treated MDA-MB-231 cells shows a small change during the experiment setting the threshold for DMSO treated cells in this experiment. (b) Specific growth rate distribution of MDA-MB-231 cells treated with .08 μM of doxorubicin has a reduced mean and a Hellinger distance from the control greater than the threshold indicating a response. (c) Specific growth rate distribution of MDA-MB-231 cells treated with .4 μM of doxorubicin has a much greater Hellinger distance from the control due to the depression of both the mean and standard deviation of the distribution.

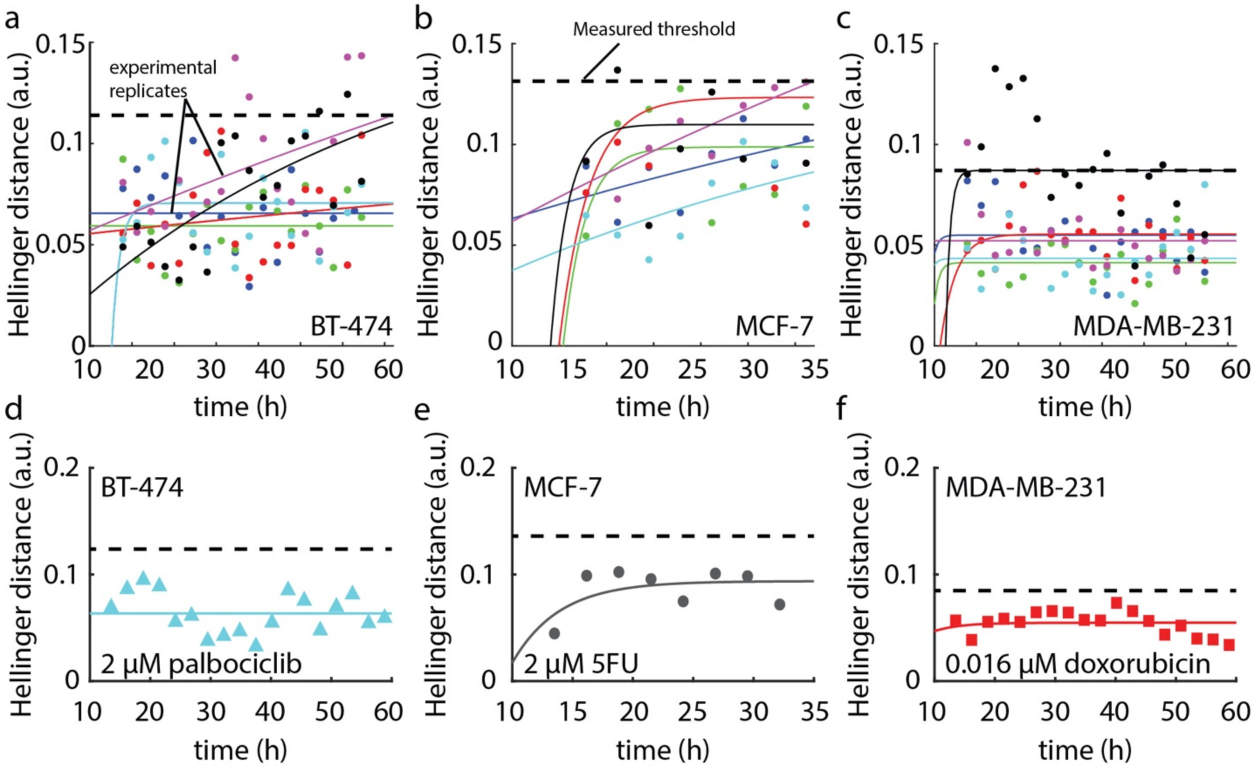


**Supplementary Figure S9: Measuring Hellinger distance.** (a) Hellinger distance over time between the DMSO treated BT-474 cells and the untreated control shows that the maximum measured Hellinger distance is .12 to determine the threshold Hellinger distance used to determine ToR. (b) Conditions that show now response never cross this threshold because the Hellinger distance between the treated group and the control is similar to the Hellinger distance between the two controls.

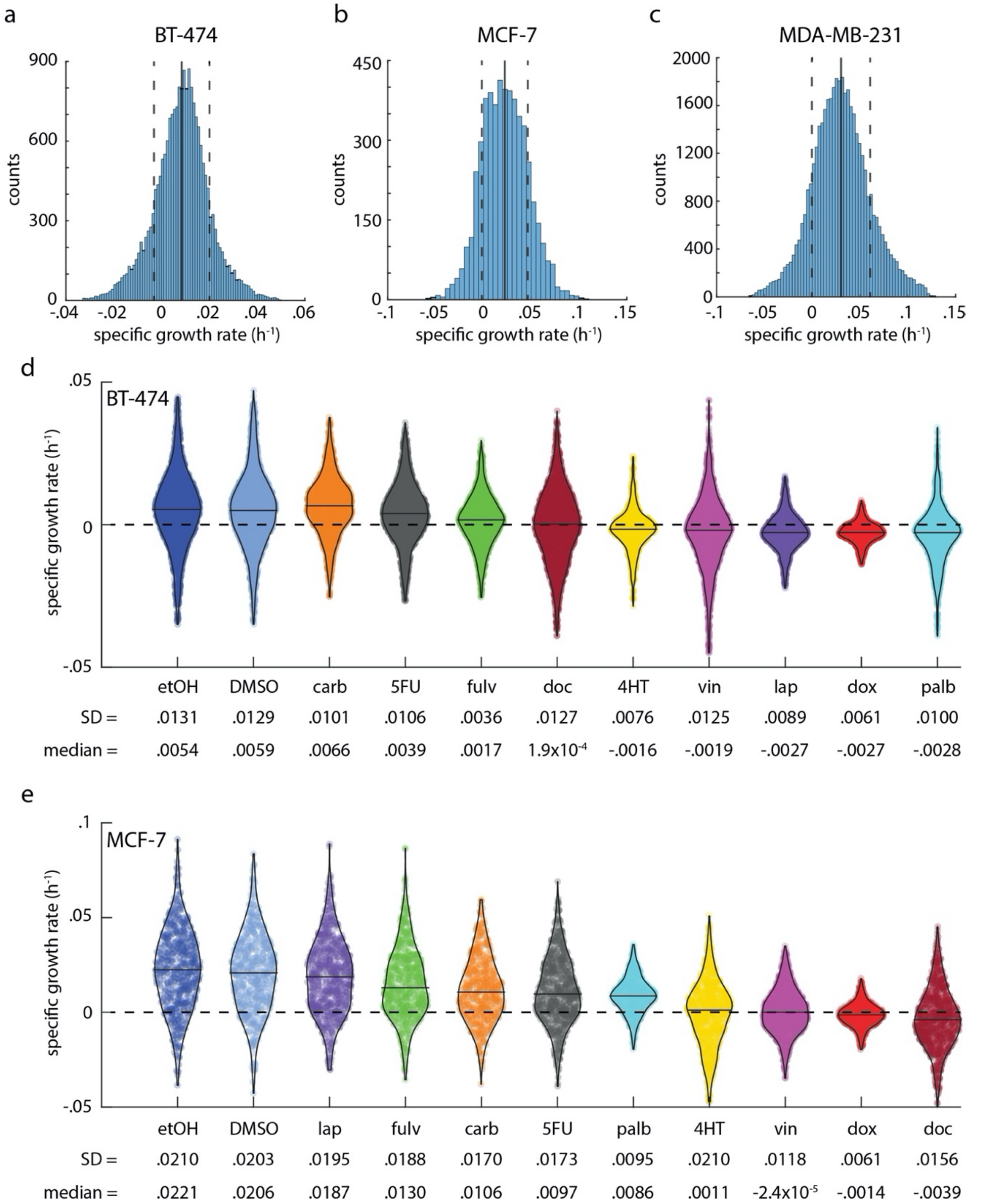


**Supplementary Figure S10: Changes in intrinsic heterogeneity during treatment are both drug and cell-line dependent.** (a-c) Each cell line exhibits baseline growth heterogeneity in the control population. Mean shown as vertical solid line, mean +/- standard deviation shown as dashed lines. (BT-474 *n* = 20,593, MCF7 *n* = 5,218, MDA-MB-231 *n* = 42,559). (d) Growth rate distribution of BT-474 cells exposed to 20 μM of each indicated drug. Data is ordered from highest to lowest mean growth rate. (e) Growth rate distribution of MCF7 clusters exposed to 20 μM drug ordered from highest to lowest mean growth rate.

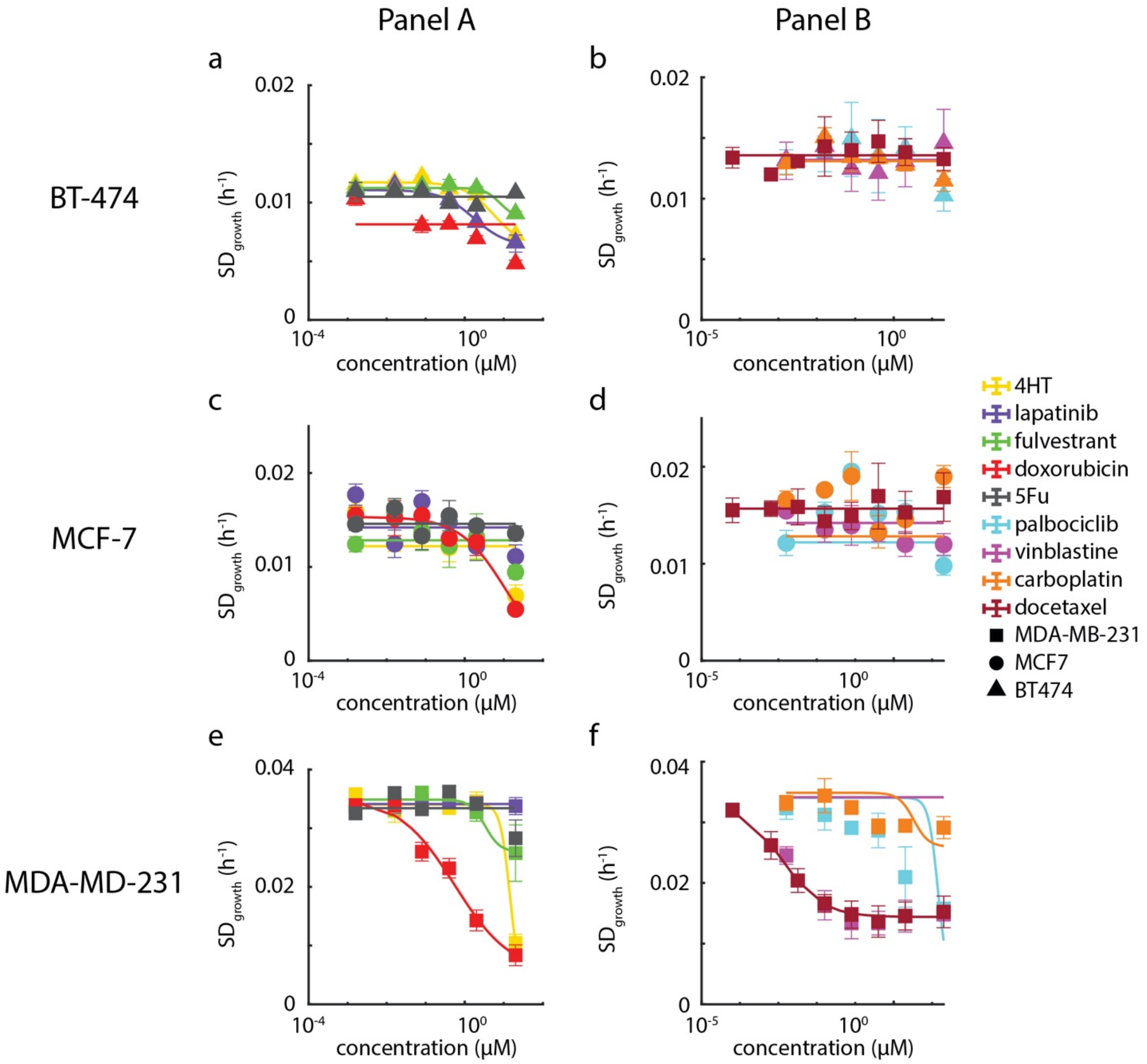


**Supplementary Figure S11: Dose response of growth heterogeneity versus therapy concentration.** Plots show standard deviation of 72 h cell cluster population specific growth rate as a function of drug concentration. Hill equation fit is only shown when there is a measurable response, as indicated when compared to a flat line fit using an F-test to reject the null hypothesis that the two fits are indistinguishable. Only a small fraction (8 out of 18 conditions) show a measurable change in population standard deviation at increasing drug concentration. Error bars show SEM.

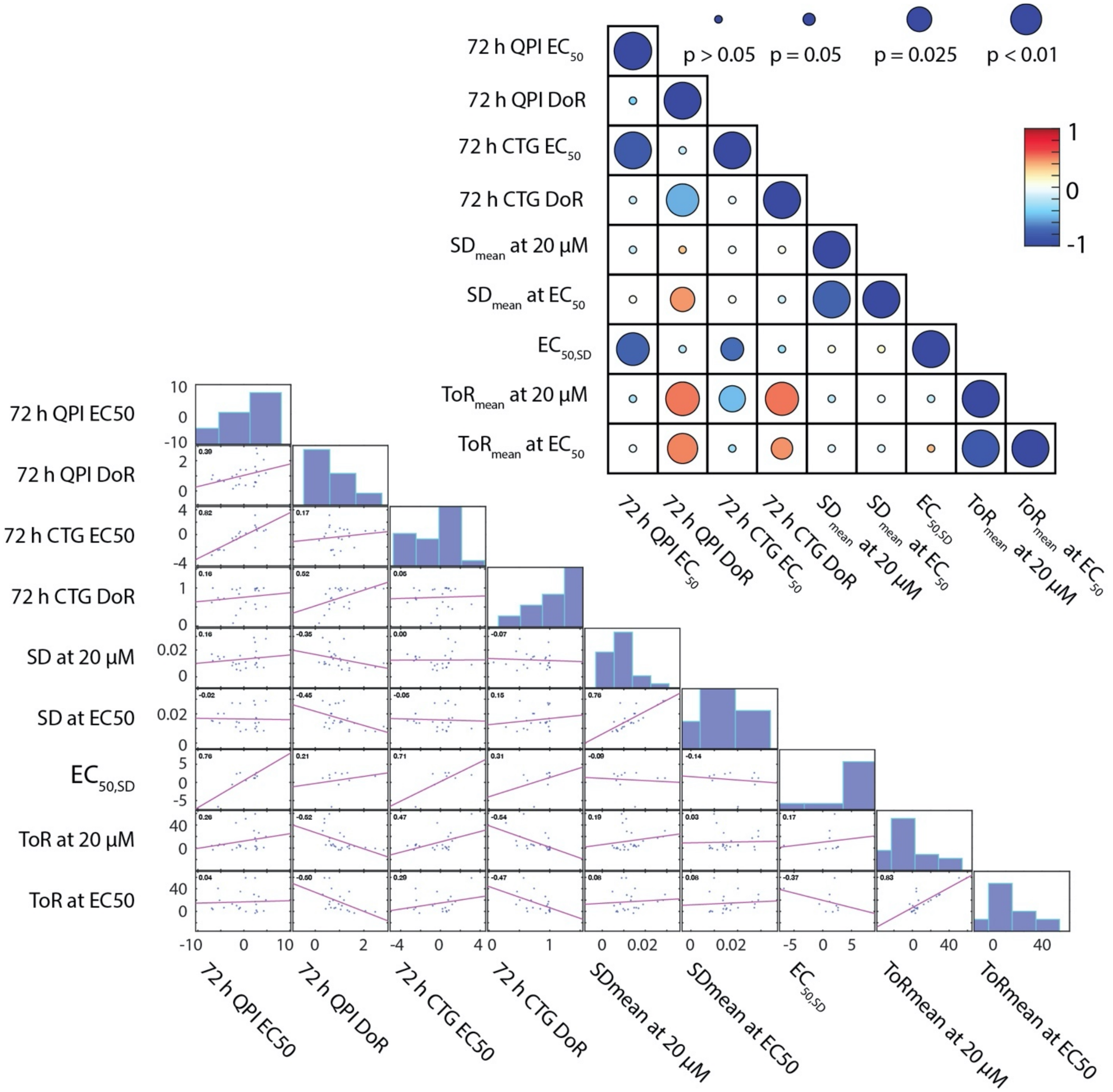


**Supplementary Figure S12: Correlations between the drug response parameters measured using QPI demonstrate orthogonality between measurements.** Correlation matrix measures the Pearson correlation coefficient and shows strong correlations as dark blue, negative correlations as red, and no correlation as white. The size of the circle shows the level of significance of the measured Pearson coefficient. Correlation plots show the data used to compute the correlation coefficient.

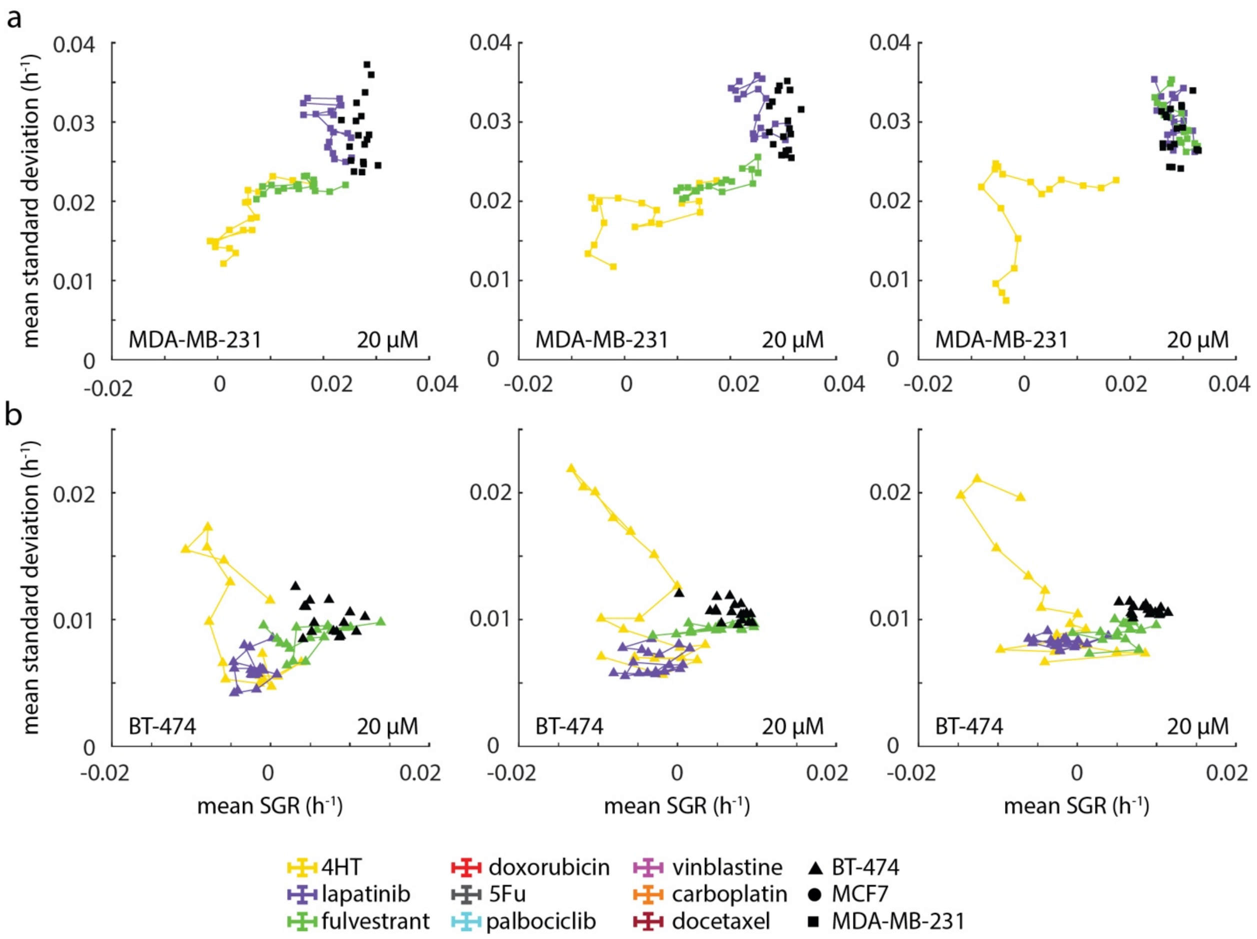


**Supplementary Figure 13: Individual replicates of time parameterized SD vs SGR.** (a) Plot of mean specific growth rate plotted against standard deviation in growth parameterized by time for MDA-MB-231 cells treated with drug panel A. Each track shows behavior from an individual experiment. (b) Plot of mean specific growth rate plotted against standard deviation in growth parameterized by time for BT-474 cells treated with drug panel A. Each track shows behavior from an individual experiment. Control cells are shown as a cluster of black points.

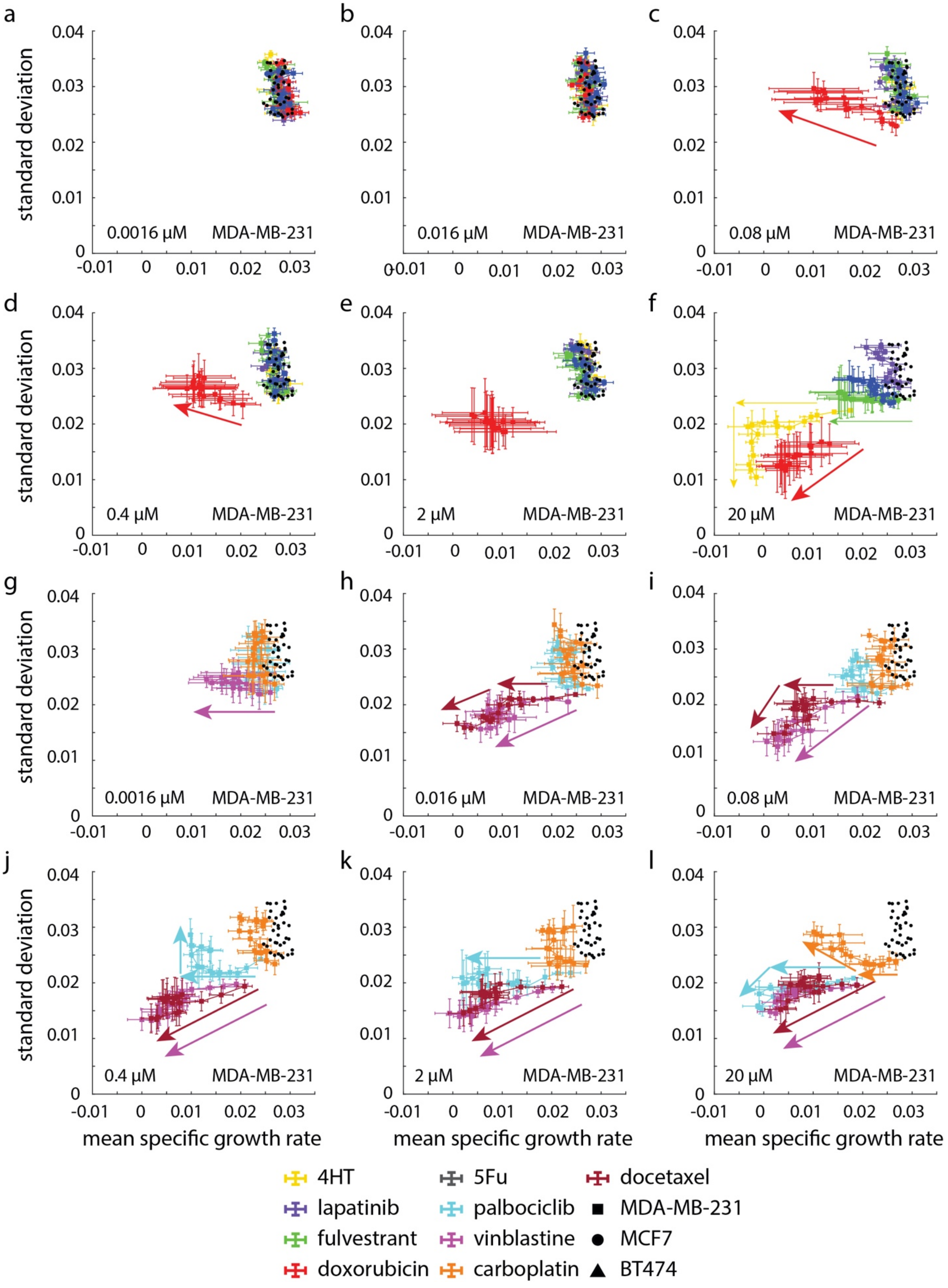

a
b
c
d
e
f
g
h
i
j
k
l
standard deviation
mean specific growth rate
0.0016 µM
0.016 µM
0.08 µM
0.4 µM
2 µM
20 µM
MDA-MB-231
4HT
lapatinib
fulvestrant
doxorubicin
5Fu
palbociclib
vinblastine
carboplatin
docetaxel
MDA-MB-231
MCF7
BT474

**Supplementary Figure S14: Heterogeneity vs specific growth rate parameterized by time for MDA-MB-231 cells shows increase in heterogeneity in response to therapy.** (a)-(b) MDA-MB-231 cells treated with drug panel A show no change in growth rate or heterogeneity over time at the lowest concentrations. At middle concentrations (c)-(e) doxorubicin shows a change in growth rate over time with little to no change in heterogeneity. (f) At the highest concentration, doxorubicin simultaneously decreases in growth rate and heterogeneity, while 4HT first decreases in growth rate and then decreases in heterogeneity. (g)-(h) At the lowest concentrations vinblastine and docetaxel start at a growth rate similar to the control and growth rate decreases over time with little change in heterogeneity. (h) At 0.08 µM MDA-MB-231 cells treated with both vinblastine and docetaxel decrease in growth rate and heterogeneity simultaneously. (j) At 0.4 µM, MDA-MB-231 cells treated with vinblastine and docetaxel decrease in growth rate and heterogeneity simultaneously, but cells treated with palbociclib first decrease in growth rate with little change in heterogeneity relative to the control. (k) At 2 µM, MDA-MB-231 cells treated with vinblastine and docetaxel decrease in growth rate and heterogeneity simultaneously, but cells treated with palbociclib decrease in growth rate with little change in heterogeneity. (l) At 20 µM, MDA-MB-231 cells treated with vinblastine and docetaxel decrease in growth rate and heterogeneity simultaneously, but cells treated with palbociclib first decrease in growth rate before decreasing slightly in heterogeneity. Carboplatin causes a decrease in growth rate with little effect on heterogeneity relative to the control. Arrows show forward direction in time. Error bars show SEM.

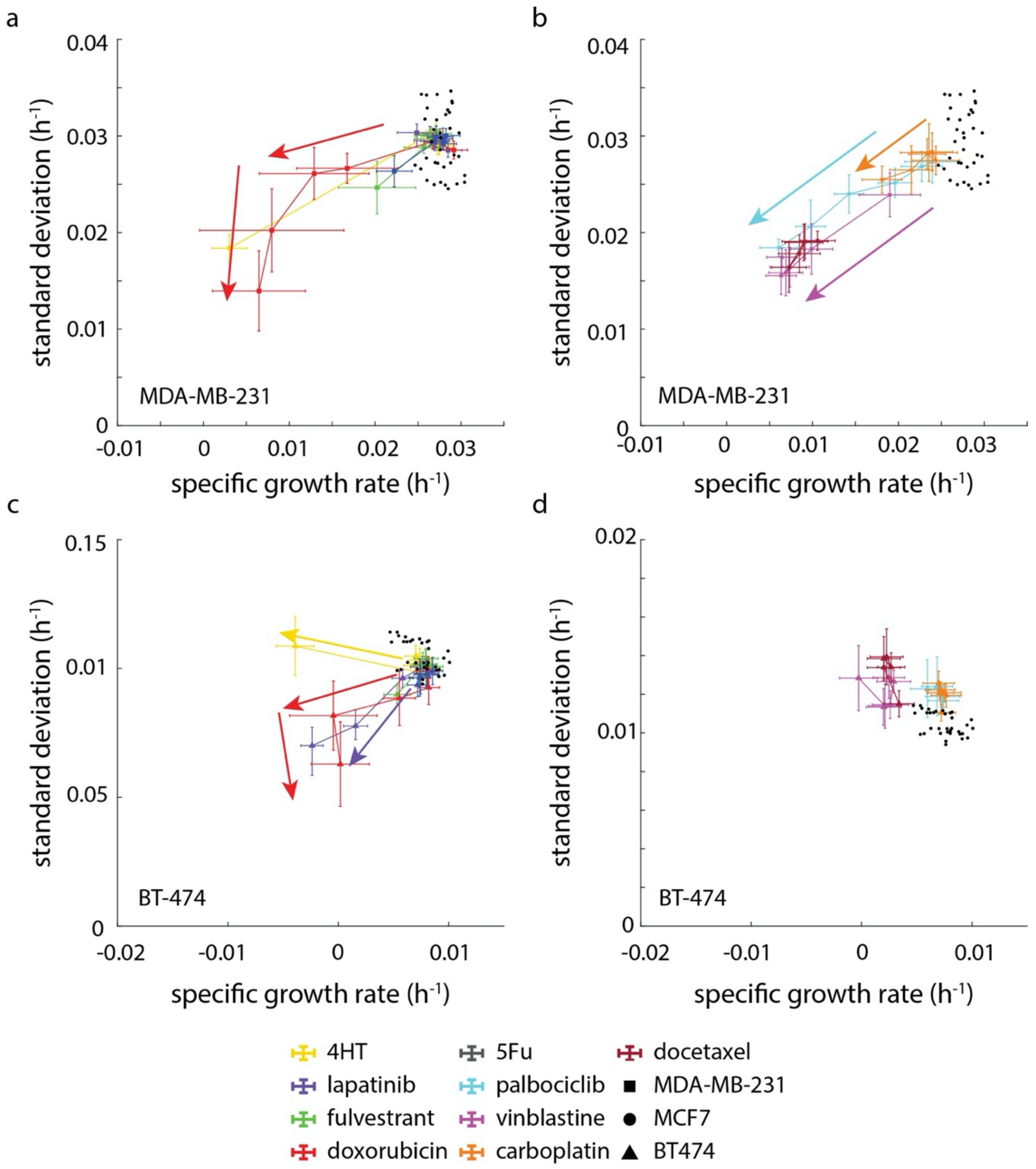


**Supplementary Figure S15: Heterogeneity vs specific growth rate parameterized by concentration for MDA-MB-231 and BT-474 cells shows increase in heterogeneity in response to therapy.** (a) As concentration increases doxorubicin treated MDA-MB-231 cells decrease in both growth rate and heterogeneity. (b) As concentration increases palbociclib, carboplatin, and vinblastine treated MDA-MB-231 cells decrease in growth rate and heterogeneity simultaneously. (c) BT-474 treated with doxorubicin, and lapatinib decrease in growth rate and heterogeneity, while 4HT treated cells decrease in growth rate at approximately constant standard deviation. (d) Doxorubicin and vinblastine treated BT-474 cells decrease in growth rate as concentration increases while heterogeneity remains constant. Arrows show direction of increasing concentration. Error bars show SEM.

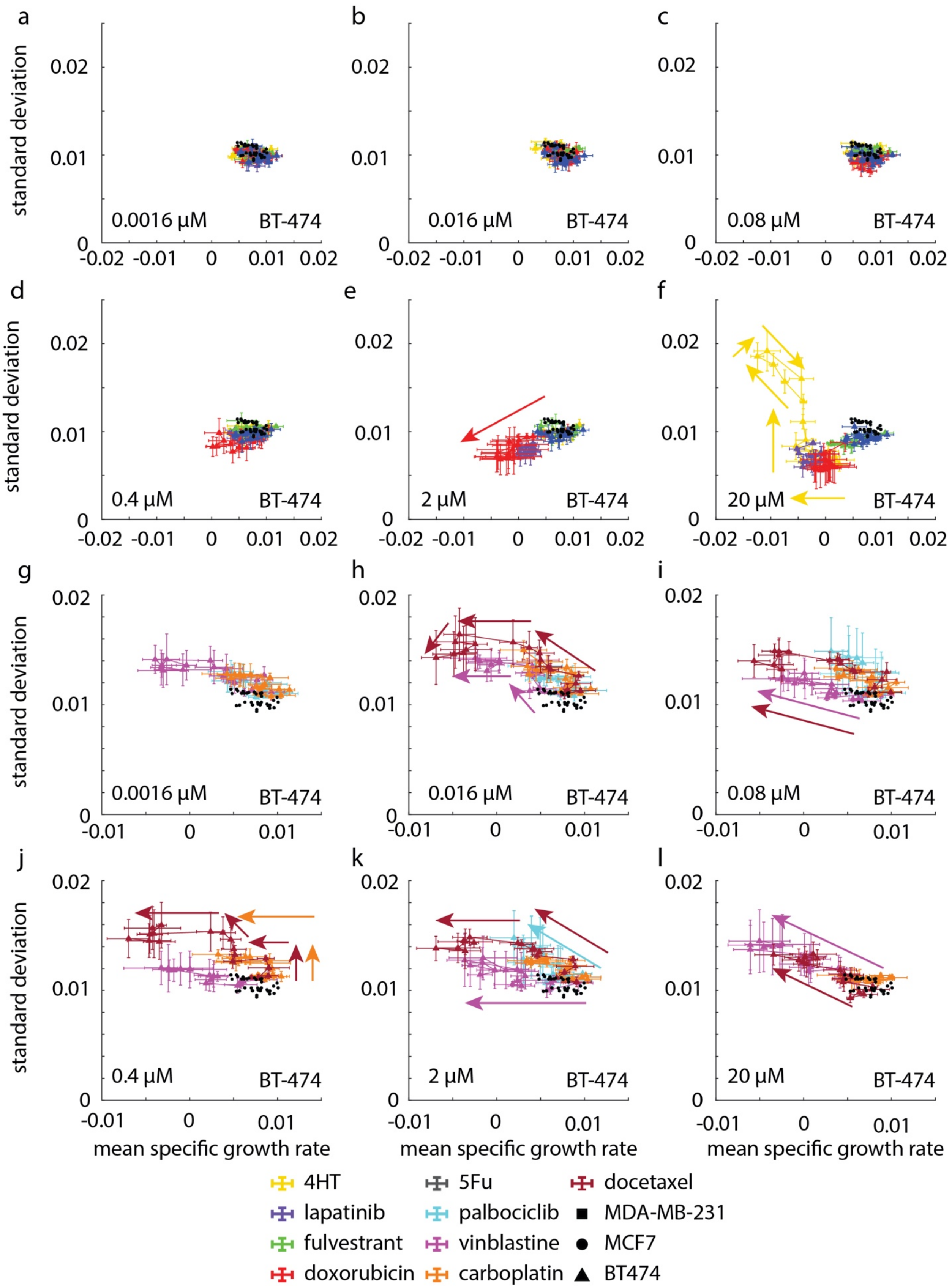

a
b
c
d
e
f
g
h
i
j
k
l
standard deviation
mean specific growth rate
0.0016 μM
0.016 μM
0.08 μM
0.4 μM
2 μM
20 μM
BT-474
4HT
lapatinib
fulvestrant
doxorubicin
5Fu
palbociclib
vinblastine
carboplatin
docetaxel
MDA-MB-231
MCF7
BT474

**Supplementary Figure S16: Heterogeneity vs specific growth rate parameterized by time for BT-474 cells shows increase in heterogeneity in response to therapy.** (a-d) At concentrations below the $EC_{50}$ both standard deviation and the mean specific growth rate for drug treated populations are indistinguishable from the control (shown as black dots). (e) Near the $EC_{50}$ concentration, the mean growth rate and heterogeneity of the population both decrease simultaneously. (f) In contrast, BT-474 cells respond to 4-hydroxy-tamoxifen by first experiencing reduced growth rate, and then a large spike in heterogeneity in response to the therapy. Only after the growth rate is further reduced does the heterogeneity of the population begin to decrease. (g) BT-474 responds to vinblastine at the lowest concentration (0.0016 μM) causing a decrease in growth rate at constant heterogeneity that is greater than the control. (h) BT-474 responds to both vinblastine and docetaxel at 0.016 μM causing a decreased growth rate and increased heterogeneity. (i) At a concentration of 0.08 μM, BT-474 cells respond with reduced growth rate and increased heterogeneity. (j) When treated with 0.4 μM of therapy, BT-474 cells respond to docetaxel with reduced growth rate and increased heterogeneity but respond to vinblastine with reduced growth rate at the same level of heterogeneity as the control. (k) At 2 μM of treatment concentration population heterogeneity is increased and growth rate decreases in response to both docetaxel and Palbociclib. (l) At 20 μM concentration the growth rate of docetaxel and vinblastine treated cells decreases, while the heterogeneity for both populations increases. Arrows show forward direction in time. Error bars show SEM.

**Supplementary movie figure captions:**

**Supplementary Movie M1:** Video of entire field of view of individual MDA-MB-231 cells treated with DMSO proliferating throughout the duration of the experiment.

**Supplementary Movie M2:** Video of entire field of view of clusters of BT-474 cells treated with DMSO proliferating throughout the duration of the experiment.

**Supplementary Movie M3:** Video of entire field of view of clusters of MCF7 cells treated with DMSO proliferating throughout the duration of the experiment.

**Supplementary Movie M4:** Video of individual MDA-MB-231 cell treated with DMSO.

**Supplementary Movie M5:** Video of a cluster of BT-474 cells treated with DMSO.

**Supplementary Movie M6:** Video of a cluster of MCF7 cells treated with DMSO.

**Supplementary Movie M7:** Video of individual MDA-MB-231 cell treated with 2 μM doxorubicin.

**Supplementary Movie M8:** Video of a cluster of BT-474 cells treated with 2 μM doxorubicin.

**Supplementary Movie M9:** Video of a cluster of MCF7 cells treated with 20 μM fulvestrant.

**Supplementary Movie M10:** Video of MDA-MB-231 cell growing despite treatment with 20 μM docetaxel.

### Description of Supplementary Data Files

**Supplementary Data 1:** All fitting parameters for all conditions and all experiments

**Supplementary Data 2:** All derived response parameters from all conditions and experiments
